# Human genomic regions with exceptionally high or low levels of population differentiation identified from 911 whole-genome sequences

**DOI:** 10.1101/005462

**Authors:** Vincenza Colonna, Qasim Ayub, Yuan Chen, Luca Pagani, Pierre Luisi, Marc Pybus, Erik Garrison, Yali Xue, Chris Tyler-Smith, The 1000 Genomes Project Consortium

## Abstract

**Background:** Population differentiation has proved to be effective for identifying loci under geographically-localized positive selection, and has the potential to identify loci subject to balancing selection. We have previously investigated the pattern of genetic differentiation among human populations at 36.8 million genomic variants to identify sites in the genome showing high frequency differences. Here, we extend this dataset to include additional variants, survey sites with low levels of differentiation, and evaluate the extent to which highly differentiated sites are likely to result from selective or other processes.

**Results:** We demonstrate that while sites of low differentiation represent sampling effects rather than balancing selection, sites showing extremely high population differentiation are enriched for positive selection events and that one half may be the result of classic selective sweeps. Among these, we rediscover known examples, where we actually identify the established functional SNP, and discover novel examples including the genes *ABCA12*, *CALD1* and *ZNF804*, which we speculate may be linked to adaptations in skin, calcium metabolism and defense, respectively.

**Conclusions:** We have identified known and many novel candidate regions for geographically restricted positive selection, and suggest several directions for further research.

## Background

The environment, acting through selective processes, is one of the main forces that shapes the genomes of species on an evolutionary timescale [1]. A population carries a certain level of genetic variation, known as ‘standing variation’; most of these variants are neutral and have no appreciable effect on fitness, but some do influence fitness. When the environment changes, natural selection acts on these non-neutral variants (and indirectly on linked variants), and over subsequent generations their frequencies may change in response. Genetic influences on fitness are typically small, and many variants usually affect the same trait, so adaptation is generally the result of small changes in frequency of multiple alleles: a ‘soft sweep’ [2]. Occasionally, a single new mutation has a major advantageous effect and can dominate the selective response, leading to a ‘hard sweep’ [2]. In addition, sexual selection provides an additional force on the genome [3], with consequences for patterns of genetic variation in the population that are difficult to distinguish from those resulting from natural selection, although hard sweeps based on the perceived attractiveness of a new observable variant may be more frequent. In other circumstances, natural selection may favor more than one allele at a particular site in the genome, for example if the heterozygote has the highest fitness, or the rarer allele is generally at an advantage, and this can lead to similar allele frequencies in different populations, sometimes maintained for long periods [4]. Thus allele frequency comparisons between different populations can potentially be informative about several forms of selection.

Humans have a worldwide distribution, but this is, evolutionarily, recent. Human populations outside Africa originated from migration out of Africa 50-70 thousand years ago, with low levels of admixture with archaic hominins, that reached all inhabitable continents by about 15 thousand years ago [5]. Our ancestors have thus been exposed to a diverse range of environments over this time period, experiencing different temperatures, altitudes, ultra-violet radiation levels, foods and pathogens; and with the development of farming and livestock domestication in some places within the last 10 thousand years, additional changes in lifestyles. The common variants in our genomes are mostly older than the out-of-Africa expansion, and thus their frequencies are potentially informative about adaptations to these environmental changes. Indeed, both general correlations between climatic variables and the frequencies of classes of functionally-related Single Nucleotide Polymorphisms (SNPs) [6], and specific examples of variants showing both frequency differences between populations and relevant functional effects have been reported. Examples of this second class are the A allele at rs1426654 within *SLC24A5* contributing to light skin color in Europeans [7], and the T allele at rs4988235 near the lactase gene (*LCT*) leading to the ability to digest lactose as an adult in Western Asia and Europe [8]. In contrast, rs1129740 at DQA1 in the HLA locus shows similar frequencies of the two alleles (coding for Cys and Tyr respectively) in many populations, linked to both increased susceptibility and protection against conditions such as hepatitis, leprosy and AIDS, possibly as a result of balancing selection [5]. Of course, apparent allele frequency patterns of these kinds are not necessarily the result of adaptation: they can also arise as a consequence of genetic drift, background selection or genotyping error [9–11], and the relative contributions of these different processes remain to be determined.

Genetic investigations of the selective forces that have shaped the human genome have mostly been based on limited genetic information: either genotyping of known variable sites using SNP arrays, or studies focused on regions of prior interest. The ability to sequence whole human genomes on a population scale has allowed analyses to be performed using more complete datasets less biased by prior expectations [12, 13]. Within the 1000 Genomes Project, we have previously identified, validated and reported SNPs with unusually high levels of population differentiation between continents, and also between populations within continents (HighD sites) [13]. These analyses revealed that there were no variants in the genome that were fixed for one allele in one continent/population and for the other allele in another continent/population; it also re-discovered the known examples mentioned above, and identified many additional highly differentiated alleles. We have also explored the functional annotations of these HighD sites, showing that they are enriched for categories such as non-synonymous variants, DNase hypersensitive sites and site-specific transcription factor binding sites, supporting the idea that they are functionally important [14].

In the current study, we have extended these analyses in three ways. First, we have included additional types of variant from the same set of samples (indel calls and large deletions from the integrated callset [13] and additional indel calls from the exome dataset); second, we have explored the properties of the most extremely low-differentiated variants (LowD sites); and third, we have further analyzed the most highly differentiated variants to investigate the extent to which they are likely to result from selection of different kinds, or other processes. In this way, we have identified many novel candidates for geographically restricted positive selection, and suggest several directions for further research.

## Results

### Identification of regions of unusually high and low differentiation among human populations

We systematically investigated the pattern of genomic differentiation among 911 individuals from the 1000 Genomes Project. We considered 36.8 million genomic variants (35,587,323 SNPs; 1,244,127 small insertion/deletions, INDELs; 13,110 large deletions, structural variants or SVs) from the 1000 Genomes Phase I integrated dataset [13] and 7,210 additional previously-unreported high quality exomic INDELs from the same samples (see Materials and Methods). We restricted our analysis to the 911 individuals belonging to the three major continental groups; samples with small sample size (IBS) and known extensive admixture (CLM, MXL, PUR) were excluded from this analysis (Supplementary Table 1).

In order to find genomic sites with high differentiation between populations, we calculated derived allele frequency differences (ΔDAF) at all variants for pairs of continents and pairs of populations within continents. This measure, ΔDAF, is highly correlated with *F*_ST_ (Spearman r 0.93 −0.95, all p-values <10e-16 in a set of 5,000 random sites from chromosome 20; Supplementary Figure 1). We performed extensive simulations to demonstrate that, like *F*_ST_, ΔDAF has >98% power to detect partial and complete selective sweeps under the range of conditions examined, and performs better than XP-EHH for partial sweeps (Supplementary Figure 2). For each pair of populations/continents, genomic ΔDAFs were ranked and a rank p-value was assigned. This analysis showed that sites with high differentiation were often strongly clustered in the genome (Figure 1), and in some cases a single differentiation event dominated the distribution (Figure 1B). Thus a list of the most highly differentiated sites based on a simple threshold would include redundant variants resulting from the same biological event, and miss other events. Assuming that one variant in each cluster drove the differentiation, and the other variants in the cluster reflect the limited recombination during differentiation, we devised the following strategy to capture the key variants. We subdivided the 1% most extremely differentiated variants into non-overlapping windows, exploring different window sizes, picked the most highly differentiated variant in each window, and retained it if it exceeded a specified ΔDAF value, again exploring a range of values (Supplementary Figure 3). As a result, we chose a window size of 5,000 variants and ΔDAF thresholds of >0.7 and >0.25 between and within continents, respectively. These choices provided a balance between stringency (enriching for the most highly differentiated variants) and number of sites (retaining sufficient for the group analyses described below). We applied a similar strategy to the exomic INDELs, but because of the smaller number of variants we only applied the ΔDAF threshold criteria without further filtering by window size. After removing variants with low genotype quality, we identified 1,137 unique variants, roughly corresponding to one HighD site every 2.5 Mb, with 248 variants (232 SNPs, 13 INDELs, 3 exomic INDELs) derived from comparisons between continents, and 903 (663 SNPs, 131 INDELs, 3 SVs, 106 exomic INDELs) from comparisons between populations within continents (Supplementary Table 2).

**Figure 1.**
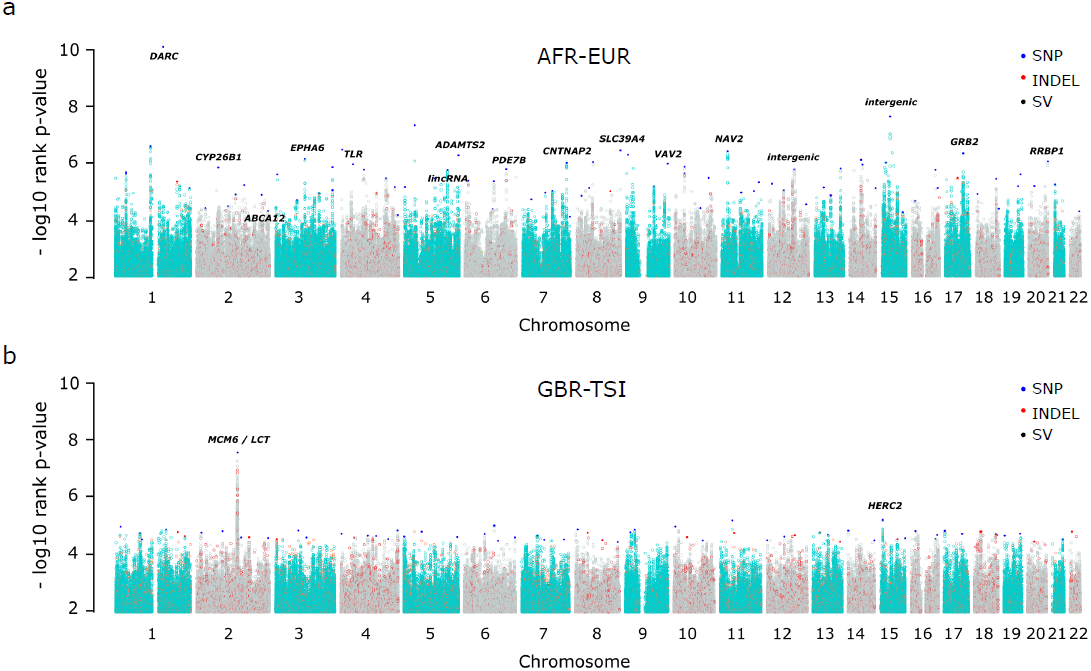
Manhattan plots of rank p-values for genome-wide pairwise derived allele frequency differences (ΔDAFs) between continental populations (a, AFR-ASN) and those within Europe (b, GBR-TSI). Only the top percentile of the 36.8 M variants is represented. Each dot represents the genome-wide ΔDAF rank p-value at a variable site between two populations. Chromosomes are represented by alternating turquoise and grey colors; red, blue and black dots represent INDEL, SNP and SV sites, respectively, that have been identified as highly differentiated (HighD sites). Gene/region names are shown for some of the top HighD sites.

We also identified regions of low differentiation among continents, and among populations within continents. In addition to the factors taken into account for HighD sites, we considered two additional features. Rare sites inevitably show low differentiation, but this is a consequence of their rarity rather than of possible balancing selection, and such sites were thus excluded. We also observed a class of sites with allele frequency 0.5 in all populations, which are likely to represent miscalled paralogous variants. For these reasons, we restricted our analysis to approximately two million sites (1,835,365 SNPs; 172,191 INDELs; 285 SVs) with 0.40 ≤ DAF ≤ 0.60 (in the entire sample of 911), excluding sites with 0.45 < DAF < 0.55. As a measure of low differentiation, we calculated the coefficient of variation of derived allele frequencies among populations (cvDAF). We ranked cvDAF genome-wide and again assigned an empirical p-value, with the lowest p-value corresponding to the smallest cvDAF. As with HighD sites, LowD sites also tended to cluster in the genome (Supplementary Figure 3) and thus we explored a range of combinations of cvDAF and window size (Supplementary Figure 4) to retain variants with the lowest cvDAF, conditioning on cvDAF <0.01 in non-overlapping windows of 100 variants. We identified 236 variants that met these criteria (213 SNPs and 23 INDELs; Supplementary Table 3).

The majority of HighD variants are SNPs (85%) and the INDEL/SNP ratio in HighD sites with good genotype quality (genotype imputation quality from MaCH/Thunder >0.8) is similar to that in matched controls (0.10 in HighD, 0.12 in controls; Fisher exact test p-value = 0.25). We validated the HighD and LowD sites with several approaches, in addition to the Sequenom genotyping reported previously [13]. First, we observed high genotype concordance at 808 sites that were independently genotyped in 112 individuals in both Phase I and Complete Genomics datasets (0.99 for one lowD INDEL. Medians: 0.98 for 629 HighD SNPs; 0.94 for 8 HighD INDELs; 0.99 for 170 lowD SNPs). Second, we compared allele frequencies between the Phase I populations and 255 independent individuals from the same populations genotyped on a different platform [15]. While we observed a high correlation of allele frequencies and ΔDAF values for HighD sites (r = 0.82-0.97, p-value <2.0e-10), the correlation of the coefficient of variation for LowD sites was poor (r = 0.13, p-value = 0.278), despite a good correlation in allele frequencies (r 0.94-0.98 and 0.77-0.96 in HighD and lowD sites, respectively; Supplementary Figure 6). Thus while both sets of genotypes are reliable and the high levels of population differentiation measured by the ΔDAF value were replicated in independent samples, the low levels of population differentiation measured by the cvDAF value were specific to the Phase I samples and are not a more general feature of the populations that the samples are derived from. We have therefore not analyzed the LowD sites further, and concentrate below on the HighD sites.

### Do HighD sites result from selection or drift?

We expect the HighD sites to result from a combination of drift and positive selection, and wished to evaluate the relative contributions of these two evolutionary forces. We therefore estimated the number of HighD sites expected under neutrality in between-continent comparisons using sequence data simulated under three established demographic models for these populations [16, 17]. We anticipated that the number of HighD sites under neutrality would be smaller than the observed number, and the difference would provide an estimate of the number resulting from non-neutral processes. In contrast, the opposite result was obtained in all two simulations: the number of simulated neutral HighD sites was higher than the observed number (Figure 2 and Supplementary Figure 8). One possible explanation for this finding could be that the neutral models were not a good fit to the Phase 1 data, and indeed we found that the number of HighD sites inferred from the simulations was extremely sensitive to parameters such as the migration rate used and the estimate of the size of the accessible genome (Figure 2, Supplementary Figure 7, Supplementary Figure 8). Since these parameters are difficult to estimate more precisely and we do not have model-independent values for them, we conclude that simulations do not currently provide a useful way to estimate the relative contributions of selection and drift to HighD site numbers. This does not necessarily indicate a general deficiency of the demographic models, but rather that they are not well-calibrated for these most extreme variants.

**Figure 2.**
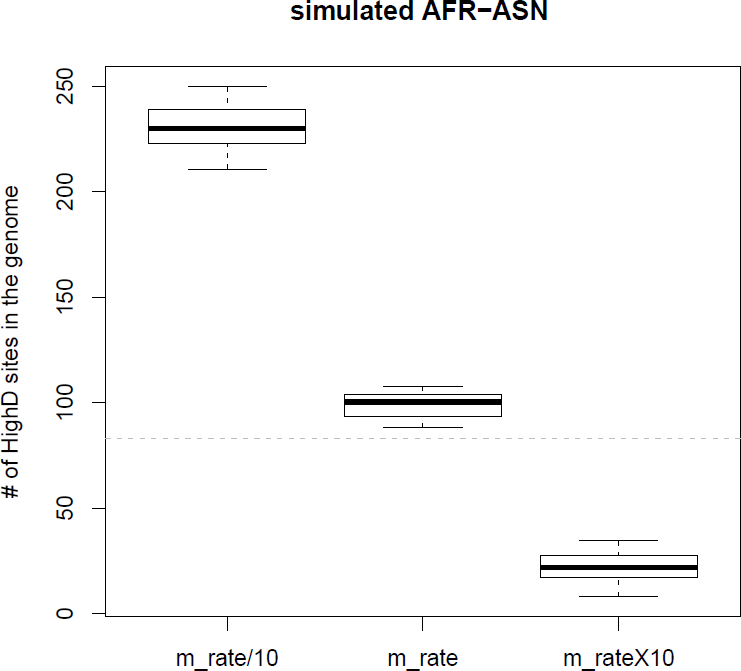
Sensitivity to migration rate (m_rate) of the expected number of HighD sites from simulations under neutrality. The dashed line represents the observed number of HighD sites. Simulations of the AFR-ASN comparison are shown here; results for AFR-EUR and ASN-EUR, and for simulations using alternative demographic models, are reported in Supplementary Figure 8.

We therefore turned to alternative sources of insights. Published studies have listed genes showing evidence of positive selection in genome-wide scans; although these lists differ between studies, a core set of genes detected by multiple scans has been compiled [18–20]. Genes harboring HighD sites were significantly enriched in this core set (34% observed *vs* 8% on average from 100 control gene sets; one sided *t*-test p-value <2.2e-16; Figure 3). Moreover, HighD genes with the highest ΔDAF values were more strongly enriched (37% for the 4th quartile), and restricting the analysis to protein-coding genes increased the overlap to 48% (average from 100 control sets of randomly selected genes is 13%; one sided *t*-test p-value <2.2e-16). The range of ΔDAF values of the HighD sites in overlapping genes was nevertheless 0.72-0.98 in between-continent comparisons and 0.25-0.63 in within-continent comparisons, suggesting that genes exhibiting the full range of ΔDAF values considered here may, according to other criteria, be positively selected, with support for selection in more than a third.

**Figure 3.**
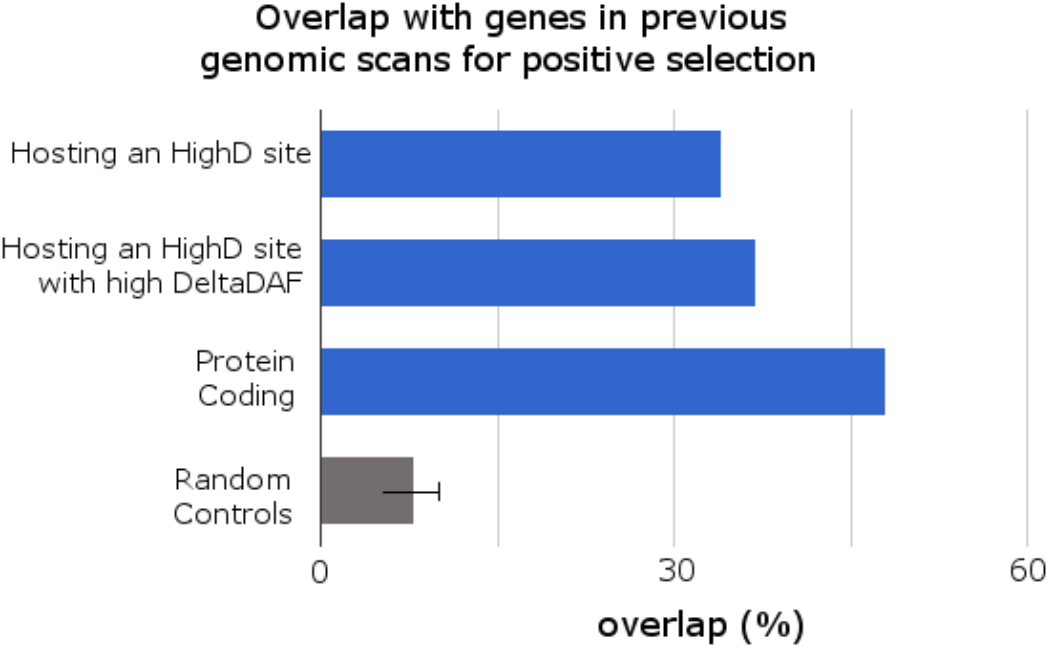
Overlap of genes hosting HighD sites with genes previously identified as putatively under positive selection. For comparison, the average value and standard deviation relative to 100 control sets of randomly-selected genes is reported. “with highDeltaDAF” refers to HighD sites whose ΔDAF is ranked in the fourth quartile.

We next looked in our own data for specific evidence of positive selection in the regions around the HighD sites (both genic and non-genic) identified through pairwise continental comparisons. We assumed that the selected allele was the derived one and that it had been selected in the population where its frequency was higher. We calculated haplotype-based statistics (iHS [21] and XP-EHH [22]) which are sensitive to recent positive selection, to assess the evidence for selection at both HighD sites and a set of randomly selected genomic variable sites matched for allele frequency and distance from genes (Supplementary Figure 9). HighD sites were found to have higher iHS and XP-EHH values (i.e. to be under selection) specifically in the population with the highest DAF (Figure 4). A similar analysis using alternative tests for selection (a combined p-value based on Tajima’s D, Fay and Wu’s H and a composite likelihood ratio test [23]) also provided support for positive selection in the population with the highest DAF, as well as for weaker selection in some additional populations (Supplementary Figure 10).

**Figure 4.**
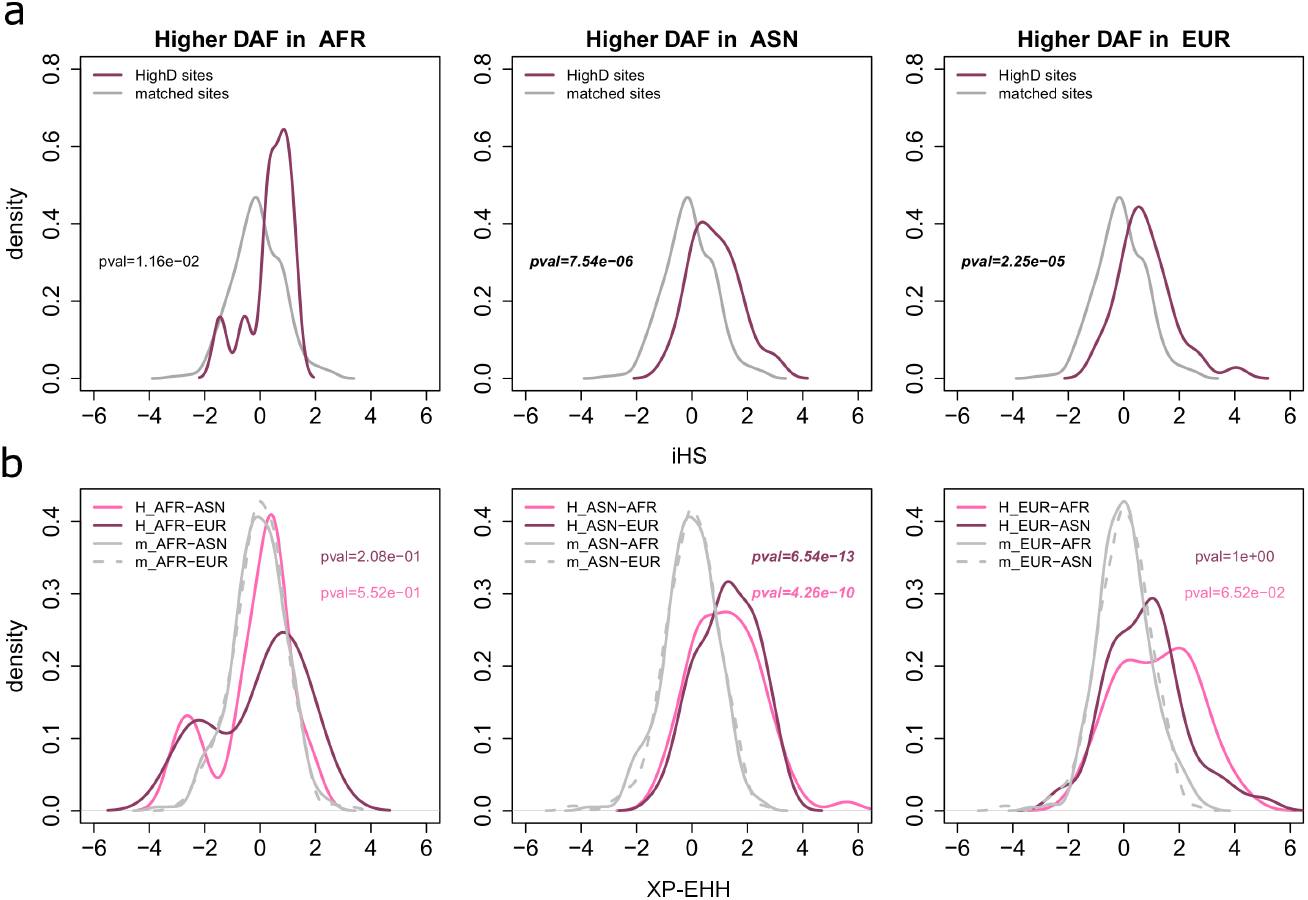
Population-specific values of iHS (4a) and XP-EHH (4b) at HighD sites. In both cases values of the two statistics relative to randomly selected genomic variable sites matched for allele frequency and distance from gene are also shown (indicated as “matched” or “m_ ”, gray lines). p-values refer to two-sample Kolmogorov-Smirnov tests between iHS or XP-EHH distributions in HighD and matched sites. In (b), for every population XP-EHH was calculated using as reference the two others (two shades of pink).

Since in a few cases specific functional targets of positive selection have been identified, we further investigated the relationship between these known targets and the variant identified by the HighD analysis. In all three examples from comparisons between continents, the precise functional target was identified correctly. These were rs1426654 in *SLC24A5* and rs16891982 in *SLC45A2*, both associated with light skin pigmentation in individuals with European ancestry [7, 24] and rs2814778 in *DARC* [25] (Figure 5a) encoding the Duffy blood group antigen O allele and associated with malaria resistance in Africans. Similarly, in comparisons between populations within continents, rs4988235 in *MCM6* and in the promoter region of LCT, responsible for lactase persistence [8] and rs12913832 in *HERC2* associated with blue eye color [26, 27] were identified correctly. We therefore conclude from these analyses that a substantial proportion of the HighD sites arose by positive selection rather than by other mechanisms, and that where it could be tested, we pick out the functional variant.

**Figure 5.**
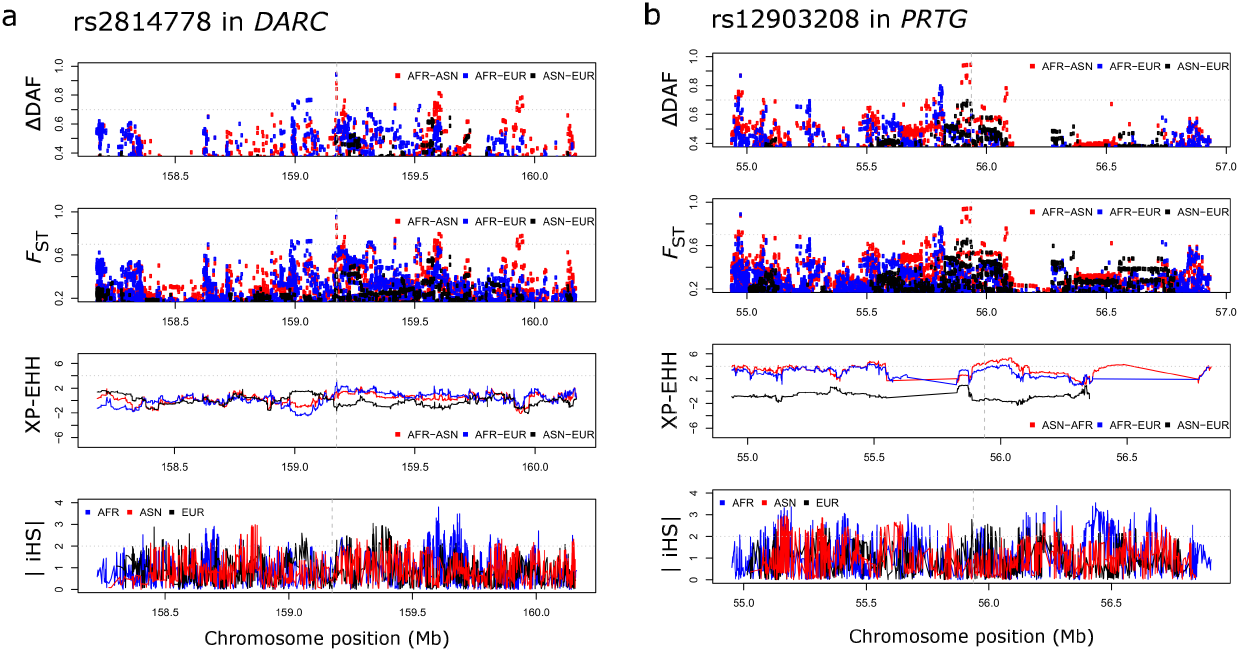
Known and novel examples of genomic regions hosting HighD sites. Comparison with other statistics informative for positive selection is also reported. Dashed vertical grey lines in the plot indicate the position indicate the position of the HighD site. In the case of DARC, this HighD site corresponds to the known functional polymorphism rs2814778. Dotted horizontal grey lines indicate reference thresholds taken from the literature for statistics for XP-EHH and iHS, or arbitrary for *F*_ST_, and as chosen in this study for ΔDAF.

### Do HighD selected sites represent cases of hard or soft sweeps?

With this knowledge, we could investigate how often positive selection in humans, ascertained in this way, acted via a classic selective sweep compared with acting on standing variation. We took two approaches to this question, first further exploring some of the relevant properties of hard sweeps by simulations, and second comparing with accepted examples of hard sweeps. We expected that a hard sweep would lead to the selected variant often lying on a single haplotype (plus its descendants resulting from mutation and recombination), while under neutrality or a soft sweep it would be more likely to lie on multiple diverse haplotype. We therefore extracted from the simulations presented above (to explore the power of the ΔDAF statistic to detect selection) the 2 kb haplotype carrying the derived allele at each selected HighD SNP, or a frequency-matched control. To provide an estimate of how different the haplotypes surrounding each target variant are from each other, we calculated the average pairwise Levenshtein distance [28] between the major haplotype and each alternative haplotype, weighted by the relative frequency of the alternative haplotype. For HighD sites resulting from a simulated selective sweep, there was often a single predominant haplotype surrounding the HighD sites (Supplementary Figure 11), and the average pairwise Levenshtein distance was 0.065 for sweeps with final allele frequency of the allele under selection between 0.8 and 1, compared with 0.126 for sweeps with final allele frequency between 0.4 and 0.6 (Supplementary Figure 12). Using as a reference the set of five HighD sites mentioned above as accepted examples of classic sweeps (i.e. *DARC*, *HERC2*, *MCM6*, *SLC24A5*, *SLC45A2*), there was always a single major haplotype (Supplementary Figure 13) and mean weighted Levenshtein distance in this reference set was 0.026. For 30 HighD sites in the empirical dataset (including that in *MCM6*) there was a single haplotype surrounding the site (Supplementary Table 4); at the other extreme, there were three cases with more than 100 haplotypes, the largest number being at rs2789823 in *VAV2* which had 185 haplotypes in the AFR population. Overall, the average weighted Levenshtein distance was 0.10, lower than in matched control sites (Supplementary Figure 14, Wilcoxon test p-value = 1.2e-4). Using as a conservative estimate for the number of HighD sites resulting from classic sweeps the number of sites with a weighted Levenshtein distance ≤0.026 (i.e. the average in the set of five reference sites), and conditioning on recombination being within the range observed at the reference sites (i.e. 0-0.025 cM/Mb), we found that 66% of the HighD sites have haplotype characteristics matching the reference set, but 36% of such sites are likely to result from chance (Supplementary Figure 15). Thus, we estimated that the true fraction of sites with features similar to reference sites is 30% (i.e. 66% minus 36%) and this is significantly more than in matched genomic regions (Supplementary Figure 15, Fisher Exact test p-value = 3.4e-2; the moderate p-value reflects the small numbers of sites in this final comparison). In Figure 6, we illustrate an example of likely selection on standing variation at the intronic HighD site rs71551254 located in the *CALN1* gene (Levenshtein distance = 0.183; DAF 0.85 in EUR, 0.05 in ASN, 0.26 in AFR), compared with a case of an apparent classic selective sweep at the intronic HighD site rs2553449 in *GSTCD* (Levenshtein distance = 0.001; DAF 0.85 in ASN 0.03 in AFR and 0.58 in EUR). In conclusion, several lines of evidence provide consistent evidence for enrichment of positive selection at HighD sites, and classic selective sweeps in at least a minority of cases.

**Figure 6.**
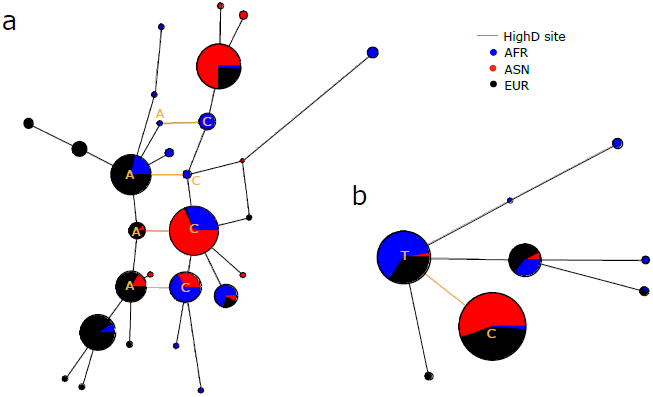
Examples of likely selection on standing variation (a, C→A at rs71551254 in *CALN1*) and a classic sweep (b, T→C at rs2553449 in *GSTCD*) inferred from high and low Levenshtein distances, respectively.

### HighD sites are enriched in genes and other functional elements

Having established the likelihood of positive selection acting on a proportion of HighD sites, probably via both hard and soft sweeps, we turned to an investigation of the probable functional targets of this selection. We first considered HighD site association with annotated genes coding for either proteins or RNAs. There was a strong enrichment for genic sites compared to frequency-matched controls (odds ratio = 6.9, Fisher exact test p-value <1e-10). We did not find enrichment for HighD sites with reported eQTLs [29], but have previously reported their enrichment in nonsynonymous SNPs and UTRs among coding sequences, and DNase hypersensitive sites and binding sites of some classes of transcription factors among non-coding regulatory regions [14]. These findings suggest that changes in both amino acid sequences and regulatory elements may have been selected.

In order to determine whether or not specific classes of genes were enriched for HighD sites, two approaches were used. Functional annotation clustering was carried out using the Database for Annotation, Visualization, and Integrated Discovery (DAVID v.6.7, [30]). Using the highest classification stringency with a Bonferroni correction for multiple comparisons, no significant enrichment of any Gene Ontology (GO) term associated with any biological process, cellular compartment, or molecular function was found. However, using medium classification stringency, a significant enrichment was observed for transcription factor binding sites at the continental and all population levels. In addition, we analyzed the same sites by Ingenuity Pathway Analysis. Here also, no significant associations were observed after applying a Bonferroni correction for multiple comparisons. Thus HighD sites, and by implication the local adaptations that underlie a proportion of them, involve diverse biological pathways, none of which predominates at our level of sensitivity.

### Novel insights into individual genomic candidates for population-or continent-specific positive selection

At the continental level, 58% (65% when restricting to protein coding) of the HighD variants are in genes previously reported to be under positive selection. Within continents, 24% (30% when restricting to protein coding) of the HighD genes belong to lists of genes putatively under positive selection. We have thus discovered a large number of HighD sites that appear to represent novel candidates for selection. Confirmation of this possibility would require functional investigations, which are beyond the scope of the current study. However, we note that while this manuscript was under review, one of our candidates, rs1871534 in *ZIP4*, was the subject of such a detailed study, and strong functional support for positive selection acting on altered zinc transport was obtained [31]. Encouraged by this, we now present a number of other candidates that illustrate a range of features.

Among the within-continent comparisons, the C→T intronic polymorphism rs77943343 is located in the caldesmon 1 (*CALD1*) gene, which encodes a calmodulin- and actin-binding protein regulating smooth muscle and non-muscle contraction. Almost half of the JPT sample carries the derived allele at rs77943343 (DAF = 0.49), which is less frequent in the other Asians (CHB = 0.15, CHS = 0.18) and continental populations in general (AFR = 0.18, EUR = 0.05). This variant lies immediately adjacent to another polymorphism, rs77994671, which, however, appears to reflect an independent mutation whose derived allele has frequency 0.024 in AFR and 0.002 in EUR but is absent in ASN. rs77943343 lies in a candidate enhancer region and in a site for histone H3 acetylation in skeletal muscle myoblasts, 497 bp downstream of an inactive promoter and 11.5 kb upstream of the active promoter (11.6 kb from the gene’s first codon) and thus might be relevant to the regulation of this gene (Supplementary Figure 16). CALD1 is implicated in Ca^++^-dependent smooth muscle contraction. Knockout of *CALD1* paralogs in zebrafish results in altered intestinal peristalsis [32] and in humans *CALD1* has been associated with gastric cancer [33] and endometriosis [34]. Network analysis of haplotypes surrounding rs77943343 in the ASN populations shows the increased frequency of a core haplotype specific to the JPT and six rare single-step derivatives also carrying the derived allele, all also present in the JPT and five specific to this population (Supplementary Figure 16). We speculate that the derived allele could have improved the efficiency of the use of Ca^++^ in conditions of low dietary calcium intake, such as the Japanese diet.

We observed two other examples of HighD sites located in genes related to calcium metabolism, both with high DAF in the LWK: rs7818866 in *VDAC3* and rs6578434 in *STIM1* (DAF = 0.67 and 0.63, respectively, compared with DAF <5% in other populations). *VDAC3* is a voltage-dependent anion channel type 3, essential for sperm mobility, while *STIM1* generates the Ca^++^ ions in oocytes during fertilization and is essential for this process. These three examples together highlight the potential importance of calcium metabolism in human adaptation.

From between-continent comparisons we present two cases. First, the intronic polymorphism rs10180970 in the ATP-binding cassette, sub-family A member 12 (*ABCA12*) has a DAF of 0.96 in Asia and 0.91 in Europe, compared with 0.13 in Africa. Haplotype network analysis of the region surrounding the HighD SNP illustrates this population differentiation and the expansion of a single haplotype cluster outside Africa, indicative of a classic selective sweep (Figure 7). ABCA12 is a member of the ABC family of proteins that transport molecules across cell membranes. Its expression is regulated by ultraviolet light [35][35] and loss-of-function mutations in *ABCA12* cause the severe skin disorder autosomal recessive congenital ichthyosis (OMIM: 607800;[36, 37]). *Abca12*(-/-) mice closely reproduce the human disease phenotype, and provide evidence that *ABCA12* plays important roles in lung surfactant and skin barrier functions [38]. Knockdown of the zebrafish ortholog similarly resulted in a phenotype including altered keratinocyte morphology [39]. Other members of the ABC family have also been identified in previous scans for positive selection [40]. The HighD SNP in *ABCA12* lies near a cluster of DNase I hypersensitive sites and a candidate enhancer specific to epithelial cells and epidermal keratinocytes (Figure 7). We suggest that this gene has been positively selected for altered expression as part of the adaptation of the skin in populations outside Africa, perhaps as a result of altered exposure to UV radiation.

**Figure 7.**
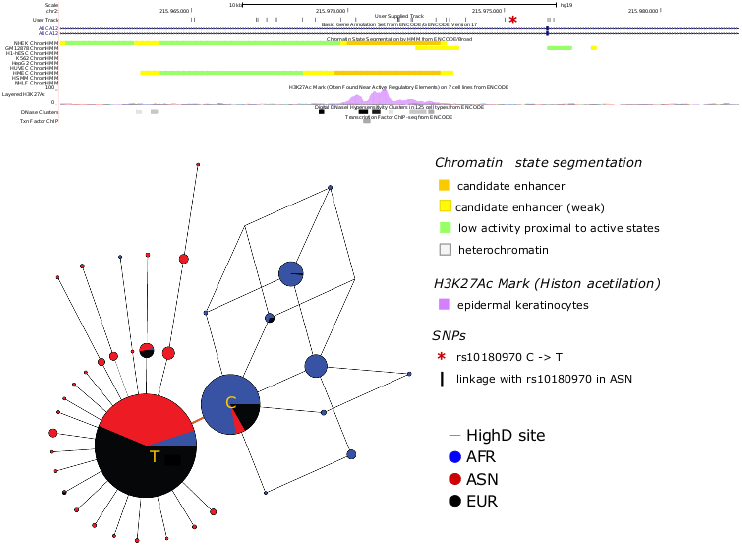
Functional annotations in the genomic region surrounding the HighD site in *ABCA12*, and a median-joining network of the haplotypes surrounding the site. Haplotypes are derived from sites in linkage disequilibrium (D’=1) with the HighD site in ASN populations.

Second, the derived allele at an exomic INDEL (a novel variant, Supplementary Table 2, chromosome 2, position 185802211) is an insertion of 3 bp (ACA) which adds a threonine residue at amino acid position 697 within the first exon of the zinc finger protein 804A gene (*ZNF804A*). This variant is present at high frequency in the ASN population (DAF = 0.83 vs 0.58 and 0.07 in EUR and AFR, respectively). ZNF804A acts as a transcription factor and regulates the transcription of genes related to schizophrenia [41]. An intronic polymorphism (rs1344706) in this gene has been associated with schizophrenia [42] and a third variant, rs4667001 in the fourth exon, changes both an amino acid and mRNA levels [43]. The ACA insertion is in strong LD with rs4667001 (r＾2 = 0.96) but less so with rs1344706 (r＾2= 0.45). Because of the threonine insertion, the protein has an additional site for post-translational modifications such as glycosylation and phosphorylation. Phosphorylation of other proteins (e.g. the deubiquitinating enzyme OTUB1) has been demonstrated to regulate susceptibility to pathogens of the *Yersinia* family [44], of which some members probably evolved in China [45], and thus we speculate that the insertion may have been selected in relation to pathogen resistance.

## Discussion

Positive selection acts through several mechanisms, and the predominant ones in human populations remain to be clarified. Population differentiation is one of the most straightforward ways to identify a subset of variants experiencing this form of selection, and indeed some of the resulting phenotypes have been recognized and studied by physical anthropologists for over a century, while the genes and specific variants that underlie them are now being identified by geneticists [5]. Whole-genome sequences from population samples provide a powerful starting resource for this, and are now available from the 1000 Genomes Project [12, 13]. In the current study, we have examined a near-complete catalog of the accessible sites in the genome that represent extreme differentiation between and within the African, European and East Asian population groups participating in the project. In doing this, we have rediscovered known examples of classic selective sweeps, and also encountered a large number of novel sites, and have thus been able to both characterize the general features of this mode of selection and obtain new insights into some specific examples.

While our validation studies confirm that the vast majority of the HighD sites we have listed are indeed highly differentiated between continents or populations, rather than being artifacts arising from genotyping errors, it is more difficult to assess the proportions that result from random genetic drift compared to positive selection. A major underlying limitation is the paucity of positively selected variants supported by direct functional evidence. HighD sites are undoubtedly enriched for positive selection (Figure 3, Figure 4), and overlap with genes previously reported as positively selected. This 34% overlap, divided into 58% for comparisons between continents and 24% for comparisons between populations, provides one estimate of the proportion that are selected, albeit one that ignores the novel events. The lower proportion in population comparisons could reflect, among other factors, the higher noise contributed by drift and the greater difficulty of detecting selection by other approaches at less complete sweeps.

This study is more comprehensive than previous surveys of highly-differentiated variants, which in large-scale studies have been limited to examining the SNPs included on genotyping arrays [11, 46] or SNPs in protein-coding genes discovered by resequencing three populations [12]. Our strategy of choosing the highest ΔDAF value from a fixed window expands the range of signals we can discover, but will nevertheless exclude weaker but still potentially interesting selection signals from the windows with a strong signal, and future studies could use more sophisticated approaches to identifying peak signals. A level of population differentiation higher than that at the *DARC* gene has sometimes been used as a criterion, and here we observe novel ΔDAF values greater than *DARC* at three positions: rs6014096 in *DOCK5*, rs1596930 in *EXOC6B* and rs12903208 in *PRTG* (Figure 5b), all in introns. In addition, genes such as *CALD1*, *ABCA12* and *ZNF804A* provide intriguing examples for follow-up in model organisms, the last also illustrating the importance of considering variants other than SNPs.

Our data throw light on two topics of current debate about selection in humans. First, some but not all studies [18] have found more evidence for recent positive selection outside Africa than inside. It has been difficult to interpret the results of tests that incorporate haplotype structure, because recombination differs between populations, with lower levels of linkage disequilibrium and different *PRDM9* alleles and recombination hotspots in Africa [47]. Similarly, SNP ascertainment in African populations has been less thorough than in European populations, and so analyses based on known SNPs have been biased against discovering highly differentiated sites in Africans. The identification of HighD sites from full sequence data, however, is unaffected by recombination or ascertainment, and the lower number of HighD sites in Africa (25 *vs* 110 each in EUR and ASN) supports the hypothesis of less positive selection of this type, despite the larger effective population size which should make selection more efficient.

Second, our identification of a large sample of likely positive selection events allows us to consider the relative importance of classic sweeps compared with selection on standing variation in humans. To do this, we need to know the proportion of HighD sites that result from positive selection rather than drift, and the proportion that result from classic sweeps. If we took the proportion of HighD sites that overlap genes with published evidence of positive selection as a minimum estimate of the proportion of selective events (58% for continental comparisons), and the proportion of inter-continental HighD sites with low Levenshtein distances (see Results) as a minimum estimate of the proportion of classic sweeps (30%), we would estimate about one-half. Both of these proportions are likely to be under-estimates – DARC would be excluded from the classic sweep numbers, for example – and thus this estimate is highly uncertain, but in contrast to some other analyses [48], emphasizes an important role for classic sweeps.

## Conclusions

This study has established a comprehensive catalog of most of the variants, including SNPs, INDELS and SVs, that are highly differentiated between the major populations of sub-Saharan Africa, Europe and East Asia. Remarkably, this simple approach, when applied to whole-genome sequences from large population samples, usually seems to lead directly to the functional variant responsible for the differentiated phenotype. Several of the most highly differentiated variants and their associated phenotypes were known long before this work, testifying to the high visibility of the phenotypes, but the remaining catalog should be a rich source of starting points for investigations of phenotypes which should be equally important and fascinating.

## Methods

We used genotype calls from two sources: the 1000 Genomes Phase I integrated callset [13] and 7,210 additional previously-unreported high quality exonic INDELs from the same samples. In each case, we restricted our analyses to 911 individuals from ten populations with ancestry from Africa, Europe or East Asia (Supplementary Table 1). Allele frequencies were calculated using vcffixup from vcflib and used to identify HighD sites. We determined distance from the nearest annotated gene using information from Ensembl v72 and subsequently generated sets of sites used in various analyses as matched random genomic sites by matching allele frequency and distance from gene start site (Supplementary Figure 10), taking into account the population to which the HighD site was assigned. Occurrence of HighD and matched sites among eQTLs was estimated using the GeneVar database [29]. Weir and Cockerham’s *F*_ST_ [49] was calculated using vcftools [50]. *F*_ST_ values were used to evaluate correlation with ΔDAF values for the same variants in chosen regions.

Using COSI, we simulated the evolution of 600 kb regions under a published demographic model [16] for three continental populations, namely AFR, EUR and ASN. We generated 3,000 replicates of neutral evolution (i.e. without any selected variant) as well as 3 × 100 replicates of 30 selective sweep scenarios as described [20], within each of the three populations. Each of these selective scenarios have a selective sweep acting on a new variant located at the center of the simulated region, i.e. at position 300,000 bp. The selective sweep lasts from 200 to 1,200 generations and always ends 401 generations ago. The selective coefficient was set to drive the selected allele up to frequency of 0.2, 0.4, 0.6, 0.8 or 1. For each replicate, we ran iHS [21], XP-EHH [22], *F*_ST_ [49] and ΔDAF [13] using a pipeline described previously [51]. For cross-population statistics, we considered all the six pairwise comparisons. We then inferred the sensitivity (true positive rate) of those four methods by (i) calculating the 95th percentile of the score distributions obtained across the 3000 neutral replicates, hence inferring the threshold corresponding to a 5% false discovery rate, and (ii) counting the proportion of replicates under a given sweep scenario with a score for the selected variant above this threshold. For each population, we performed the sensitivity analysis for five different sweep scenarios by grouping the replicates where the final allele frequency of the selected variant was the same, as well an “overall” sweep scenario grouping all the replicates with selection in a given population.

Genotype concordance of HighD and LowD sites between Phase I data and publicly available Complete Genomics data was calculated as the proportion of concordant calls among 112 overlapping individuals averaged across loci. Allele frequency concordance was calculated between Phase I EUR, AFR, ASN populations and three non-overlapping sets of individuals from the same continents from the HapMap3 study [15] (total of 255 individuals; 24 EUR: 18 CEU, 6 TSI; 166 ASN: 101 CHD, 31 JPT, 34 CHB; 65 AFR: 22 LWK, 43 YRI) genotyped on a different platform (Illumina BeadArray genotypes at 1,456,587 sites [15]). Because overlap of sites on the chip with HighD and LowD sites was poor (100 HighD and 29 LowD sites) we also included comparisons at sites in high LD (r＾2 > 0.8) with HighD and LowD sites (255 and 202 sites, respectively). LD was calculated using Vcftools [50]. Concordance was assessed using a Spearman correlation coefficient.

In order to estimate the expectation for number of HighD sites under neutrality, we simulated 100 replicates of 500 Mb DNA sequence data (subdivided into 10 chromosomes of the same size) from 911 diploid genomes from three populations according to two published demographic models [16, 17] for the three continental populations AFR, ASN and EUR using MaCS [52]; we included variable recombination rates by incorporating information from random regions from HapMap Phase3 recombination maps [15]. Because all models gave led to similar conclusions we only report results from one [17]. We estimated the number of HighD sites in 500 Mb of simulated data as for the Phase 1 data, and then scaled this number to the size of the accessible genome (2,526,390,487 base pairs [13]).

We summarized from the literature a list of 3,467 genes that have been previously identified in in genomic scans for positive selection [18–20] and compiled the occurrence of non-redundant genes hosting HighD sites (HighD-genes; n = 542) in it. To obtain estimates of random expectation we calculated the occurrence of 100 sets of control genes in the list of positively selected genes. Control genes were randomly selected to match the number of HighD-genes or fractions of this set based on ΔDAF quartiles (n = 247 and n = 146 for 3^rd^ and 4th quartiles, respectively). A one-sided t-test was used to assess the significance of the differences in overlap.

We estimated genome-wide statistics informative about selection from sequence data of the Phase I data set. These statistics were based on 10 kb windows and include three allele frequency spectrum-based tests, Tajima’s D, Fay and Wu’s H, and Nielsen et al.’s composite likelihood ratio (CLR), calculated and combined to give a single p-value as described previously [23]. We also estimated the haplotype-based statistics iHS [21] and XP-EHH [22] using Selscan [53] at sites for which recombination maps and ancestral allele information were available. In order to make the continent and population level analyses comparable, in each group we restricted the analysis to a set of 30 randomly-chosen individuals. The iHS and XP-EHH scores obtained for each SNP in each population/continent were divided into allele frequency classes and, within each class, normalized following standard procedures [21, 40]. Similarly, we calculated iHS and XP-EHH for a set of genomic sizes matched to HighD sites for allele frequency in the combined sample and distance from gene start site. A two-sample Kolmogorov-Smirnov test was used to evaluate differences between the HighD and matched site iHS distributions.

The Database for Annotation, Visualization, and Integrated Discovery (DAVID v.6.7) was used for functional characterization of the highly differentiated variants that lay within genes. The Functional Annotation Clustering option was used by adding Panther and Reactome to Pathways, Panther to Protein Domains and Reactome and UCSC TFBS to Protein Interactions to the default settings. In another approach, Ingenuity Pathway Analysis (IPA, Ingenuity Systems, Redwood City, CA) was carried out for the same set of variants. Ensembl gene identifiers were uploaded and core analyses was carried out for the differentiated variants at the continental (CON) and population levels (AFR, ASN and EUR). The analyses generated networks based on their connectivity in the Ingenuity Knowledge Base (IKB), which includes experimental data from human, mouse and rat models. The core analyses selected only those interactions which have been experimentally observed and included pathways based on 1) Diseases and disorders, 2) Molecular and cellular functions and 3) Physiological system development and function. The significance of the association between the dataset and the pathways was tabulated by estimating a ratio between the number of genes from the dataset that met the expression value cut off that map to the pathway, and the total number of molecules present in the pathway. A conservative Bonferroni p-value threshold was used to account for multiple testing.

Median-Joining networks for haplotype visualization and analysis were generated using Network 4.6.1.1 [54]. Since this version of the software can display only 100 chromosomes per circle, we selected each time 300 random chromosomes. The *CALD1* and *ABCA12* haplotypes were based on all the SNPs in D’=1 within 10 kb of the HighD site in the population with the highest DAF; for the networks in Figure 7 and Supplementary Figure 17, haplotypes were derived from polymorphisms within 2 kb surrounding the HighD site.

## List of relevant urls

The 1000 Genomes Project http://www.1000genomes.org/

vcflib https://github.com/ekg/vcflib/#vcffixup

Complete Genomics http://www.completegenomics.com/

HapMap SNP-chip data http://hapmap.ncbi.nlm.nih.gov/downloads/index.html.en#release

HapMap recombination maps http://hapmap.ncbi.nlm.nih.gov/downloads/recombination/

Ingenuity Pathway Analysis www.ingenuity.com

## Authors’ contributions

VC participated in the project design and coordination, performed analyses and drafted the manuscript. QA participated in the study design and performed analyses. YC, LP, PL, MP, EG and YX performed analyses. CTS participated in its design and coordination and helped to draft the manuscript.

## Description of additional data files

Additional files include additional figures (Colonna_20140503_add_fig.pdf), additional tables (Colonna_20140503_add_tab.xlsx), and a full list of The 1000 Genomes Project participants (The_1000_Genomes_Consortium_participants_list.pdf)

## Acknowledgements

We thank Danny Challis, Fuli Yu, Donna Muzny and Richard Gibbs for contributing the exomic indel data set and Goo Jun for helping with the Complete Genomics data set. We thank Tsun-Po Yang for assistance with Genevar. This work was supported by The Wellcome Trust (098051), an Italian National Research Council (CNR) short term mobility fellowship from the 2013 program to VC, and an EMBO Short Term Fellowship ASTF 324-2010 to VC.

